# Biochemical characterizations of ER-resident peroxidases, GPx7 and GPx8, reveal their distinct and regulated oxidative activities

**DOI:** 10.1101/2020.03.26.010116

**Authors:** Shingo Kanemura, Elza Firdiani Sofia, Naoya Hirai, Masaki Okumura, Hiroshi Kadokura, Kenji Inaba

## Abstract

In the mammalian endoplasmic reticulum (ER), the diverse network comprising more than 20 members of the protein disulfide isomerase (PDI) family and more than five PDI oxidases has evolved to promote oxidative protein folding. While the canonical disulfide bond formation pathway constituted by Ero1α and PDI has been well studied so far, mechanistic and physiological bases of newly identified PDI oxidases, glutathione peroxidase-7 (GPx7) and -8 (GPx8), are only poorly understood. We here demonstrated that human GPx7 has much higher reactivity with H_2_O_2_ than human GPx8, leading to efficient PDI oxidation. GPx7 forms a catalytic tetrad at the redox active site to react with H_2_O_2_ efficiently and stabilize a resultantly generated sulfenylated species. While it was previously postulated that the GPx7 catalysis involved a highly reactive peroxidatic cysteine, a resolving cysteine was found to act to regulate the PDI oxidation activity of GPx7. The present study also revealed that GPx7 formed complexes preferentially with PDI and P5 in H_2_O_2_-treated cells. Altogether, human GPx7 functions as an H_2_O_2_-dependent PDI oxidase in cells whereas PDI oxidation may not be the central physiological role of human GPx8.

## Itroduction

Most secretory and cell-surface membrane proteins undergo oxidative protein folding in the endoplasmic reticulum (ER) to acquire native conformations via disulfide bond formation and isomerization. In the mammalian ER, the reactions are catalyzed by a variety of oxidoreductases which encompass more than 20 members of the protein disulfide isomerase (PDI) family (1–5) and more than five PDI oxidases including ER oxidoreductin-1α (Ero1α) (6–9) and peroxiredoxin-4 (Prx4) (10–12). Some PDI family enzymes also engage in disulfide bond reduction to promote protein folding (13), the ER-associated degradation of misfolded proteins (14–16), or the ER-to-cytosol retrograde translocation of a bacterial cholera toxin and non-enveloped viruses (17–19). PDI family members typically possess thioredoxin (Trx)-like domains with a CXXC motif at the redox-active site, thereby exerting thiol-disulfide exchange reactions with substrate proteins (1,3). Thus, the PDI family enzymes, in combination with their upstream oxidases or reductases, constitute a diverse thiol-mediated network to maintain the proteostasis in the ER (2,3).

Ero1α and PDI play central role in the canonical pathway of disulfide bond formation at the cost of one molecular oxygen per disulfide bond, resulting in the generation of H_2_O_2_ as a byproduct (2,6). Overproduction of H_2_O_2_ as a consequence of the Ero1α overwork causes oxidative stress in the ER, eventually leading to the cell death. Therefore, the oxidative activity of Ero1α needs to be tightly controlled (20). In the mechanism of operation of mammalian Ero1α, four cysteine residues in the electron shuttle loop act also as components of a regulatory switch which ensures strict regulation over Ero1α activity (8,9,20–23). An ER-resident peroxiredoxin, Prx4, metabolizes H_2_O_2_ and oxidizes PDI family enzymes upon return to the reduced state (24,25). Our prior *in vitro* studies showed that ERp46 and P5, with their partner oxidase Prx4, are engaged in the rapid but promiscuous disulfide bond introduction in the initial phase of oxidative protein folding (10), whereas PDI, in concert with Ero1α, efficiently catalyzes the correction of non-native disulfide bonds and the subsequent selective formation of native disulfide bonds (26). Other oxidative pathways involving Vitamin K epoxide reductase (VKOR) were also reported to potentially operate in the PDI reoxidation cycle (27,28).

More recently, glutathione peroxidase-7 (GPx7) and -8 (GPx8) were identified as ER-resident PDI oxidases using H_2_O_2_ as a source of oxidative power (29–31). GPx-family was historically named after its firstly identified member that catalyzes the reaction of peroxide with reduced glutathione (GSH) on its selenocysteine (Sec) residue (32). Despite the absence of a GSH-binding motif and the substitution of the Sec residue by Cys in GPx7 and GPx8, both the enzymes are capable of acting as peroxidases which reduce H_2_O_2_ to water (29,31,33). GPx7-deficient cells were found to accumulate endogenous reactive oxygen species (ROS), lowering cellular viability (34,35). Another study demonstrated that GPx7 plays a role in the alleviation of oxidative stress in breast cancer cells, oesophageal cells, and adipose tissue, emphasizing the importance of GPx7 in the oxidative stress response (35–38). Both GPx7 and GPx8 were suggested to interact with Ero1α *in vivo* and possibly modulate the peroxide-generating activity of Ero1α by scavenging H_2_O_2_ in its proximity (29). Additionally, the Ero1α-GPx8 complex may serve to prevent the diffusion of H_2_O_2_ from the ER (39).

GPx7 and GPx8 commonly possess an essential peroxidatic cysteine residue (C_P_) in the typical motif, NVASXC(or U)G (Cys57 in GPx7 and Cys79 in GPx8), and a resolving cysteine residue (C_R_) in the highly conserved motif of GPx family, FPCNQF (Cys86 in GPx7 and Cys108 in GPx8) (40,41). Whereas GPx7 and GPx8 share a highly similar overall structure, their topologies are distinct in the following points: GPx7 is purely a luminal protein whereas GPx8 has a small N-terminal cytoplasmic domain, a short transmembrane domain, and a catalytic-active domain facing the lumen (named ‘luminal domain’ in this work) (33). Although some structural and molecular insights have been gained into GPx7 and GPx8, the mechanistic details of GPx7/8-mediated PDI oxidation and their physiological roles in cells, especially their involvement in oxidative protein folding, still remain unclear.

In this study, we investigated the mechanistic basis of human GPx7- and GPx8-catalyzed PDI oxidation and identified their preferential substrates among PDI family members both *in vitro* and *in vivo*. We thus demonstrated that GPx7 was a much more efficient PDI oxidase than GPx8 due to even higher susceptibility of C_P_ to H_2_O_2_, in which Gln92 contained in the catalytic tetrad plays a critical role. While the GPx7 catalysis was known to involve the C_P_ residue (31), we found that the C_R_ residue acts to regulate the PDI oxidation activity of GPx7. Our extensive studies also revealed that GPx7 bound PDI and P5 in H_2_O_2_-treated cells. Thus, human GPx7 is a potent PDI oxidase whereas the role of human GPx8 seems largely distinct from that of GPx7 despite their structural similarity.

## Results

### Higher reactivity with H_2_O_2_ and PDI of GPx7 than that of GPx8

Although both GPx7 and GPx8 are presumed to work as a PDI-dependent peroxidase (29), the reactivity of these two enzymes with H_2_O_2_ have not been assessed precisely. To compare their peroxidase activities, we first mixed the fully reduced form of GPx7 or the luminal domain of GPx8 (1 μM each) with different concentrations (10, 50 and 200 μM) of H_2_O_2_ and chased their redox state changes (Fig. 1). The aliquot samples were treated with 1 mM maleimidyl PEG-2000 (mal-PEG 2k) for quenching at indicated time points. While the topmost and bottom bands are identifiable as reduced and oxidized species, respectively, the intermediate band is most likely a hyperoxidized species (SO_2_H or SO_3_H form), in which C_R_ is mal-PEG modifiable but C_P_ is no longer alkylated due to the hyperoxidation with H_2_O_2_. In support of this, the intermediate bands significantly increased at higher H_2_O_2_ concentrations as observed particularly with GPx8 (Fig. 1, right).

**FIGURE 1.**
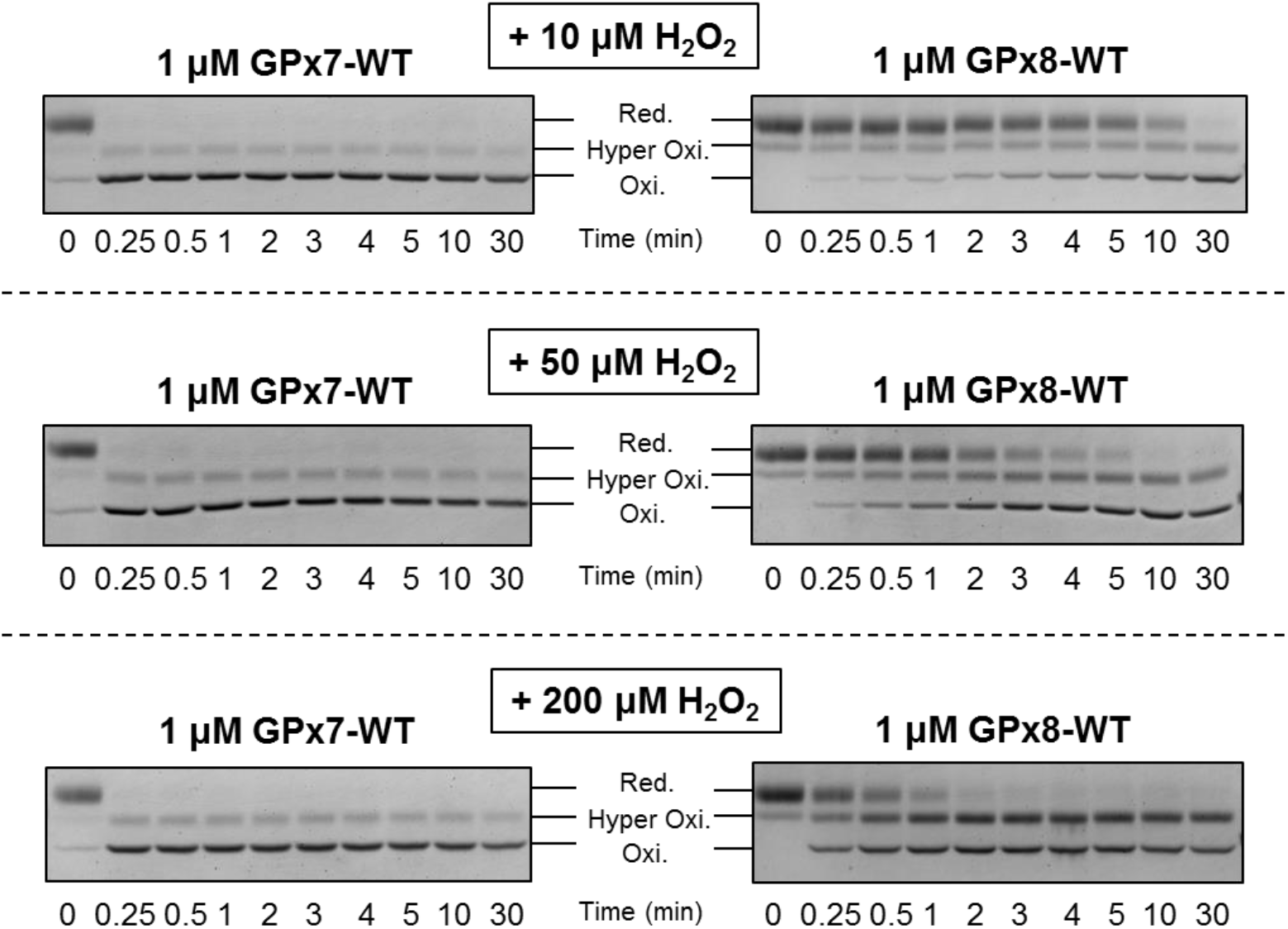
Reactivities of GPx7 and GPx8 with hydrogen peroxide. Time course of the redox-state change of GPx7 and the luminal domain of GPx8 during a reaction between 1 μM GPx7/8 and 10, 50 or 200 μM H_2_O_2_. At indicated time points, the reaction mixture was quenched with mal-PEG 2k. GPx7 and the luminal domain of GPx8 were separated by non-reducing SDS-PAGE and stained with CBB. Red., Hyper Oxi., and Oxi denote fully reduced, hyperoxidized and fully oxidized species of GPx7/8, respectively.

The present assay revealed much higher reactivity or affinity of GPx7 for H_2_O_2_ than that of GPx8; 10 μM H_2_O_2_ was sufficient to fully oxidize 1 μM GPx7 within 15 sec after mixing, whereas most fraction of 1 μM GPx8 was kept reduced even at mins of the reaction time in the presence of 10 μM H_2_O_2_. Higher concentrations of H_2_O_2_ significantly accelerated GPx8 oxidation in concomitant with increased generation of the hyperoxidized species, suggesting that the *K*_d_ for H_2_O_2_ value of GPx8 is much above 10 μM.

To visualize H_2_O_2_-driven PDI oxidation by GPx7 and the luminal domain of GPx8, reduced PDI (10 μM) was reacted with GPx7 or GPx8 (1 μM each) in the presence of 10, 50 or 200 μM of H_2_O_2_, and the redox-state change of PDI was monitored by mal-PEG 2K modification of free cysteines followed by non-reducing SDS-PAGE (Fig. 2A). Consequently, GPx7 generated significant amounts of partially and fully oxidized PDI at early time points whereas GPx8 hardly accelerated PDI oxidation compared to H_2_O_2_ alone. This result is in line with the present observation that GPx7 was oxidized by H_2_O_2_ much more efficiently than GPx8 (Fig. 1). Of note, GPx7-mediated PDI oxidation was slower at 200 μM H_2_O_2_ than at 50 μM H_2_O_2_. This is likely explained by the greater generation of a hyperoxidized GPx7 species during the catalysis of PDI oxidation in the presence of higher concentrations of H_2_O_2_. In support of this notion, most portion of GPx7 was converted to a hyperoxidized species during the catalysis of PDI oxidation at 200 μM H_2_O_2_ (Fig. 2B).

**FIGURE 2.**
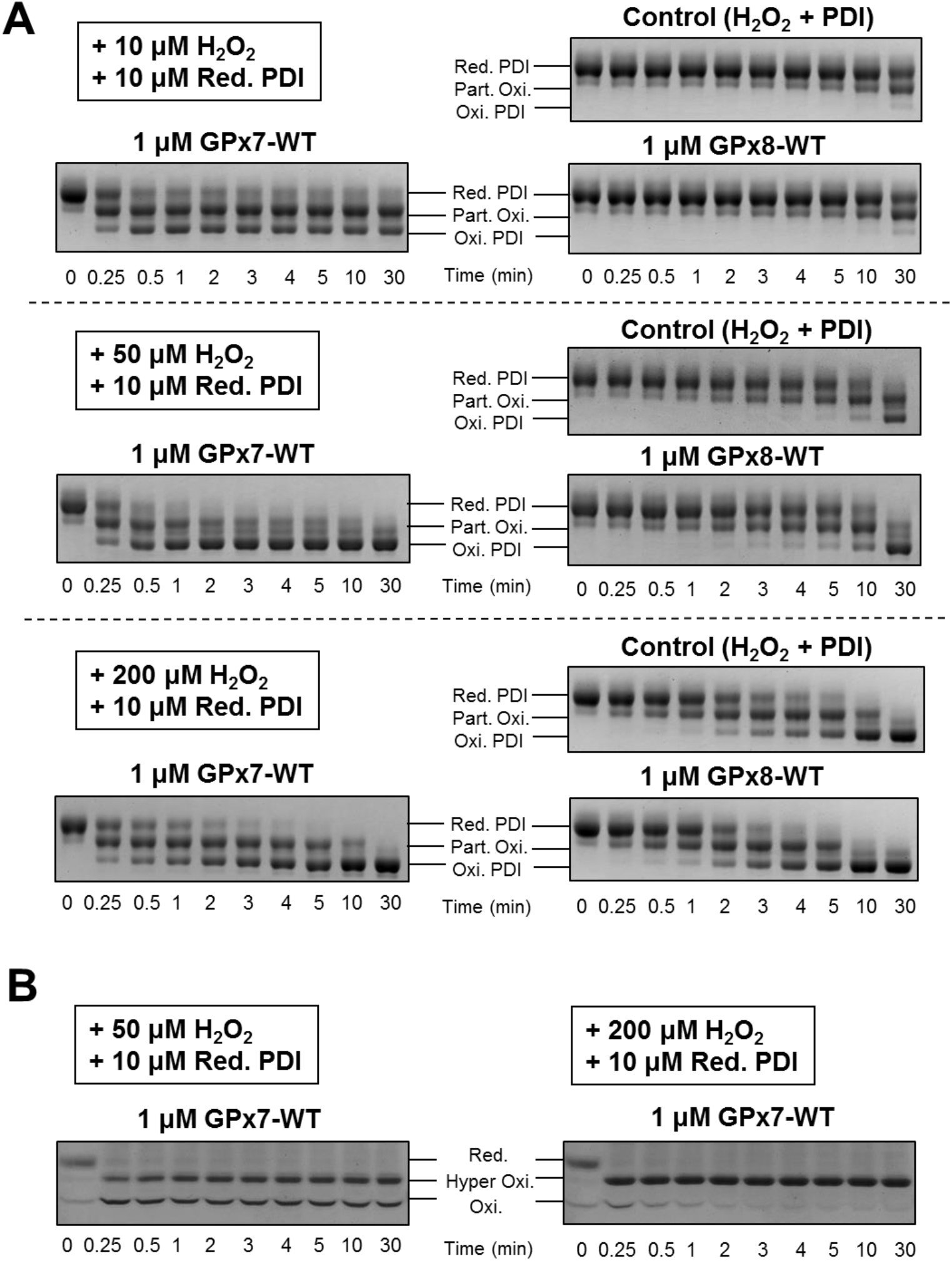
GPx7/GPx8-catalyzed oxidation of PDI in the presence of hydrogen peroxide. **A.** Time course of the redox-state changes of PDI during non-catalyzed, GPx7-, and GPx8-catalyzed PDI oxidation. All experiments were initiated by mixing 10, 50 and 200 μM H_2_O_2_ into the mixture of reduced PDI (10 μM) and GPx7 or the luminal domain of GPx8 (1 μM) in buffer containing 50 mM Tris/HCl (pH 7.4) and 300 mM NaCl. At indicated time points, the reaction mixture was alkylated with mal-PEG 2k. The redox states of PDI was separated by non-reducing SDS-PAGE and stained with CBB. Red. PDI, Part. Oxi., and Oxi. PDI denote fully reduced, partially oxidized and fully oxidized species of PDI, respectively. **B.** Time course of the redox-state change of GPx7 WT (1 μM) during the catalysis of oxidation of PDI (10 μM) in the presence of 50 or 200 μM H_2_O_2_

A p*K*_a_ value, which determines the degree of protonation at a given pH, represents a fundamental character of protein thiol groups (42,43). To examine whether the p*K*_a_ values of GPx7 and GPx8 can explain their different peroxidatic activities, we analyzed crystal structures of the enzymes (PDB ID: 2P31 for GPx7 and 3CYN for GPx8) using the program PROPKA3. The *in silico* analysis yielded the p*K*_a_ values of GPx7 and GPx8 peroxidatic cysteines (C_P_) to be 9.3 and 11.5, respectively. Thus, C_P_ in GPx8 is predicted to have exceptionally low reactivity compared with free cysteine in solution (p*K*a: 8.3) and C_P_ in GPx7. Altogether, the results tempt us to suspect a functional role of GPx8 as a H_2_O_2_-scavenging peroxidase in the ER.

### Essential role of a Gln residue in the catalytic tetrad of GPx7

Effective thiol-based peroxidases commonly contain a catalytic triad or tetrad including a highly reactive, C_P_ (40,41). Indeed, GPx7 contains a catalytic tetrad consisting of Cys57, Trp142, Asn143, and Gln92 at the active site, where Gln92 is predicted to stabilize thiolated or sulfenylated Cys57 through a hydrogen bond upon reaction with H_2_O_2_ (40). While GPx8 also conserves the first three residues, Cys79, Trp164 and Asn165, at the positions corresponding to Cys57, Trp142 and Asn143 in GPx7, respectively, we surmised that the replacement of Gln92 (in GPx7) to Ser114 (in GPx8) (Fig. 3A & B) was a primary reason for much lower peroxidase activity of GPx8 than that of GPx7. In this context, the distance between S_γ_ atom of GPx7 Cys57 and O_ε1_ of Gln92 is 3.1 Å, close enough to stabilize the thiolated GPx7 C_P_ via a hydrogen bond to the O_ε1_; meanwhile the S_γ_ atom of GPx8 Cys79 is separated from O_γ_ atom of Ser114 by 4.8 Å, far beyond a hydrogen bonding distance (Fig. 3B).

**FIGURE 3.**
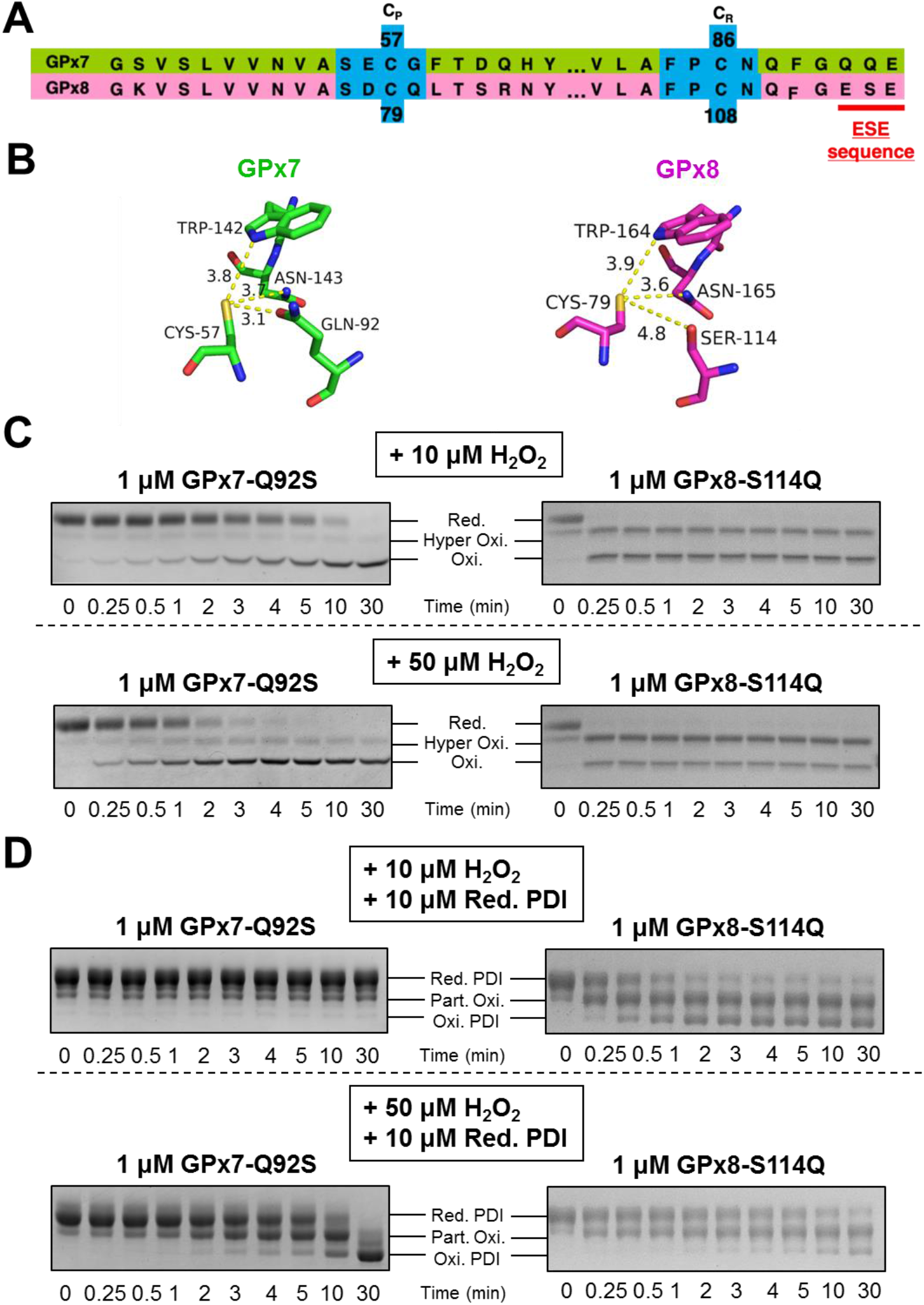
Effects of a mutation at the catalytic tetrad on the reactivity of GPx7/8 with hydrogen peroxide and PDI. **A.** Amino acid sequence alignment in the C_P_- and C_R_-neighboring regions of human GPx7 and GPx8. **B.** Catalytic tetrads of GPx7 and GPx8 involving a peroxidatic cysteine and its neighboring Trp and Gln/Ser residues. In GPx7, the O_ε1_ atom of the Gln92 side chain is distant from the S_γ_ atom of Cys57 by 3.1 Å, whereas the O_γ_ atom of Ser114 in GPx8 side chain is separated from the S_γ_ atom of Cys79 by 4.8 Å. **C.** Time course of the redox-state change of a GPx7 and GPx8 swap mutants (GPx7 Q92S and GPx8 S114Q) during a reaction between 1 μM GPx7/8 and 10 or 50 μM H_2_O_2_. At indicated time points, the reaction mixture was alkylated with mal-PEG 2k. **D.** Time course of the redox-state change of PDI during a reaction between 1 μM GPx7 swap mutant (GPx7 Q92S) or GPx8 swap mutant (GPx8 S114Q) and 10 μM PDI in the presence of 10 or 50 μM H_2_O_2_.

To explore the importance of the C_P_-neighboring residues in the different peroxidatic activities of GPx7 and GPx8, we constructed swap mutants where Gln is mutated to Ser in GPx7 (GPx7 Q92S) and vice versa in the luminal domain of GPx8 (GPx8 S114Q). In the result, the GPx7 swap mutation greatly compromised the GPx7 reactivity with H_2_O_2_ (Fig. 3C, left), leading to the abolishment of PDI oxidation activity (Fig. 3D, left). Notably, GPx7 Q92S exhibited a similar rate in H_2_O_2_-dependent oxidation as GPx8 though the latter generated larger amount of hyperoxidized species than the former at both 10 μM and 50 μM H_2_O_2_ (Fig. 3C left and Fig. 1). It is also noteworthy that GPx8 S114Q greatly increased the GPx8 reactivity with H_2_O_2_, leading to more efficient PDI oxidation in the presence of 10 μM H_2_O_2_ (Fig. 3C & D, upper right). However, GPx8 S114Q only partially oxidized PDI at 50 μM H_2_O_2_ (Fig. 3D, lower right), likely due to the predominant formation of the hyperoxidized form as observed for GPx7 WT in the presence of 200 μM H_2_O_2_ and 10 μM reduced PDI (Fig. 2B, right). Even without reduced PDI, GPx8 S114Q indeed generated a considerable amount of hyperoxidized species upon reaction with 50 μM H_2_O_2_ (Fig. 3C, lower right). Collectively, we conclude that Q92 acts as an essential residue for high reactivity of GPx7 with H_2_O_2_ and hence effective PDI oxidation of the enzyme.

### GPx7 Cys86 plays a critical role in the regulated oxidative activity of GPx7

Three-dimensional structures of reduced forms of GPx7 and GPx8 have high similarity, in which the locations and orientations of C_P_ and C_R_ in GPx7 are almost identical to those in GPx8 (29). The S_γ_ atoms of C_P_ and C_R_ are separated by approximately 11 Å in both the enzymes, suggesting that a large conformational change is required for formation of an intramolecular disulfide bond between these two cysteines (30). Given the situations, two alternative mechanisms have been proposed for GPx7/GPx8-mediated PDI oxidation: a 1-Cys mechanism in which the C_P_ acts as the sole redox center that reacts with both H_2_O_2_ and PDI, and a 2-Cys mechanism in which an intramolecular disulfide bond between C_P_ and C_R_ is formed preceding the oxidation of PDI (30,33).

To investigate which of the mechanisms is primarily exerted by GPx7 and GPx8, we constructed single-Cys mutants of the enzymes in which C_R_ was mutated to alanine: C86A for GPx7 and C108A for GPx8. Of interest, the results demonstrated that GPx7 C86A oxidized PDI more efficiently than GPx7 WT (Fig. 4A, left), suggesting that a 1-Cys mechanism is more competent in PDI oxidation than a 2-Cys mechanism and that GPx7 WT exerts primarily a 2-Cys mechanism or both 1-Cys and 2-Cys. By contrast, GPx8 C108A oxidized PDI even less efficiently than GPx8 WT (Fig. 4A, right), indicating the necessity of C_R_ for the GPx8-mediated PDI oxidation. Thus, GPx8 is likely to utilize a 2-Cys mechanism exclusively.

**FIGURE 4.**
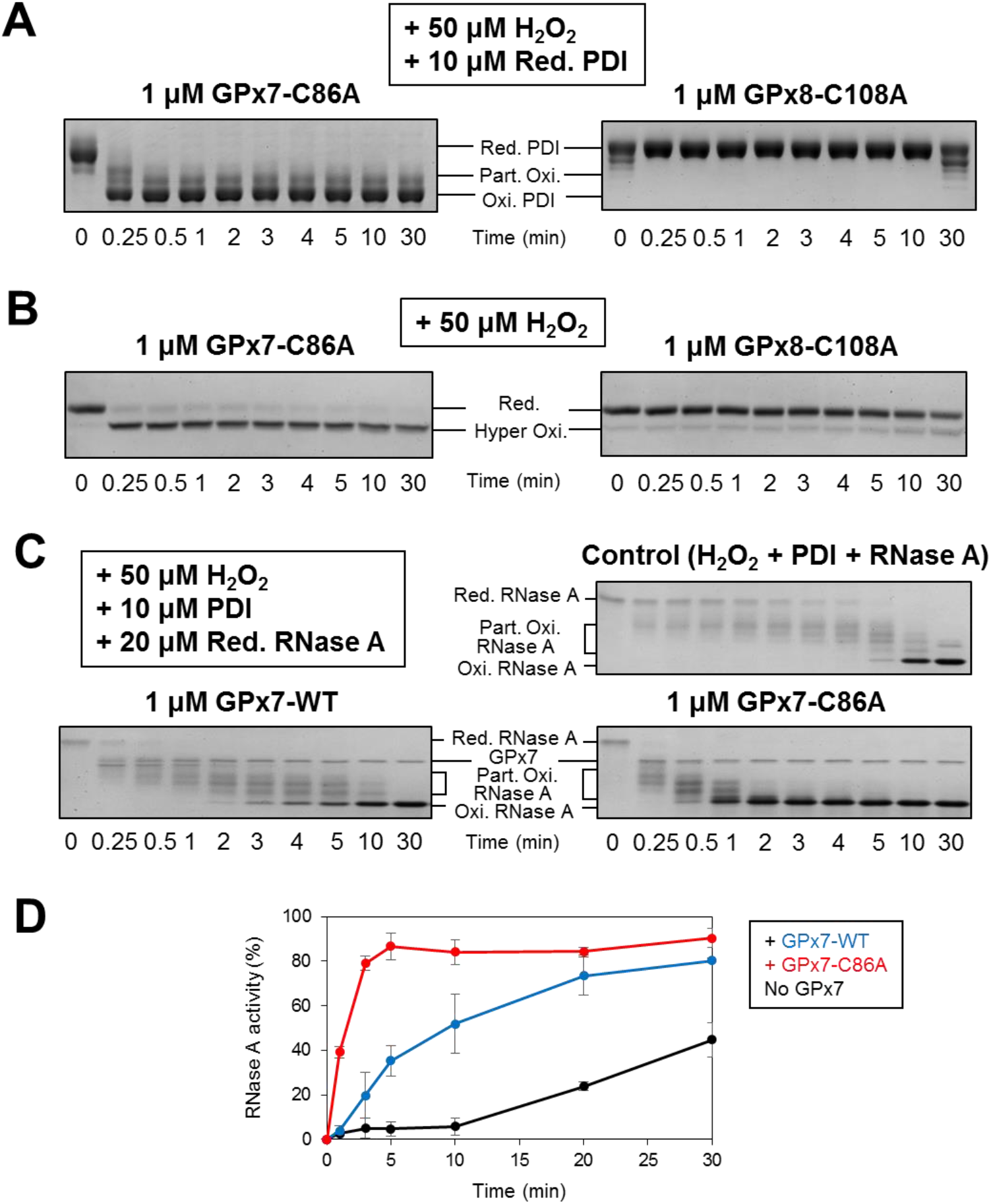
Effects of a C_R_-residue mutation on the oxidative activities of GPx7/8. **A.** Time course of the redox-state change of PDI during a reaction between 1 μM GPx7 C86A or GPx8 C108A and 10 μM reduced PDI in the presence of 50 μM H_2_O_2_. At indicated time points, the reaction mixture was alkylated with mal-PEG 2k. **B.** Time course of the redox-state change of GPx7 C86A and GPx8 C108A after mixture with 50 μM H_2_O_2_. At indicated time points, the reaction mixture was alkylated with mal-PEG 2k. Red., and Hyper Oxi. denote fully reduced and hyperoxidized species of the GPx7/8 mutants, respectively. **C.** Time course of the redox state changes of reduced/denatured RNase A (20 μM) during oxidation by 10 μM PDI and 1 μM GPx7 WT or C86A in the presence of 50 μM H_2_O_2_. At indicated time points, the reaction mixture was alkylated with AMS. Red. RNase A, Part. Oxi. RNase A, and Oxi. RNase A denote fully reduced, partially oxidized and fully oxidized species of RNase A, respectively. **D.** Recovery of RNase A activity during oxidative folding catalyzed by 10 μM PDI and 1 μM GPx7 WT or C86A in the presence of 50 μM H_2_O_2_.

For GPx7 to exert a 1-Cys mechanism with high efficiency, its C_P_ must be very reactive with H_2_O_2_. To verify that it is indeed the case, we investigated the susceptibility of GPx7 C86A to H_2_O_2_-mediated hyperoxidation. As expected, GPx7 C86A was rapidly hyperoxidized with 50 μM H_2_O_2_ whereas its counterpart, GPx8 C108A, was much less sensitive to H_2_O_2_ (Fig. 4B). To further compare the oxidative activities of GPx7 WT and C86A, we investigated oxidative folding of RNase A catalyzed by these two using gel-shift assay (Fig. 4C). The results demonstrated that GPx7 C86A oxidized RNase A much more rapidly than GPx7 WT. Accordingly, GPx7 C86A restored RNase A activity more quickly than GPx7 WT (Fig. 4D). Thus, GPx7 can be a more competent H_2_O_2_-dependent oxidase by exerting a 1-Cys mechanism exclusively on the highly reactive peroxidatic cysteine.

To explore whether GPx7 WT primarily employs a 1-Cys or 2-Cys mechanism, we examined the PDI concentration dependence of the rate of PDI oxidation by GPx7 WT and C86A by performing the NADPH consumption assay. We predicted that, at higher PDI concentrations, a 1-Cys mechanism gets more predominant over a 2-Cys mechanism since, under such conditions, reduced PDI should more likely win the race of nucleophilic attack to a sulphenylated C_P_ against the C_R_ residue of the same GPx7 molecule (Fig. 5A). Given that, the oxidative activity of GPx7 WT will approach that of GPx7 C86A as the concentration of PDI increases. However, higher PDI concentrations did not render the initial NADPH consumption rates of GPx7 WT and C86A closer to each other (Fig. 5B & C). It is thus conceivable that, regardless of the PDI concentration, GPx7 WT primarily employs a 2-Cys mechanism, resulting in the slower oxidation of PDI and hence the slower oxidative folding of a downstream substrate than GPx7 C86A (Fig. 4).

**FIGURE 5.**
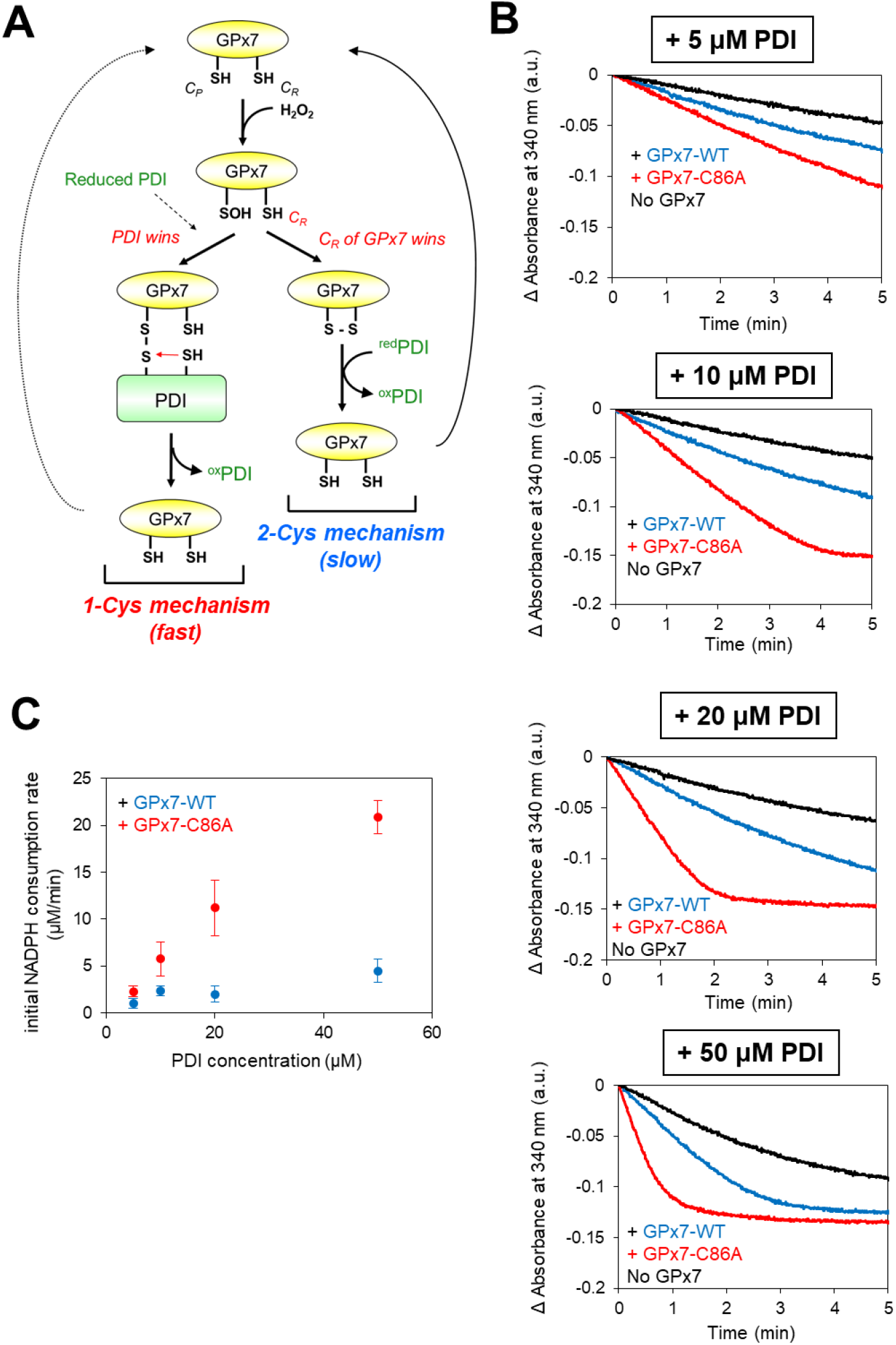
Effects of PDI concentration on the oxidative activities of GPx7 WT and C86A. **A.** Alternative (1-Cys or 2-Cys) mechanism of GPx7 in the catalysis of PDI oxidation. A peroxidatic cysteine (C_P_) of GPx7 is readily sulfenylated upon reaction with H_2_O_2_, followed by the race between reduced PDI and a resolving cysteine (C_R_) of the same GPx7 molecule against the sulfenylated C_P_. **B.** NADPH consumption coupled to PDI oxidation catalyzed by GPx7 WT or GPx7 C86A was monitored by measuring absorbance change at 340 nm. All experiments were initiated at 30°C by mixing 50 μM H_2_O_2_ into the mixture of reduced PDI (5-50 μM), GSH (1 mM), GR (1 U), NADPH (200 μM) and GPx7 WT or GPx7 C86A (1 μM) in buffer containing 50 mM Tris/HCl (pH 7.4) and 300 mM NaCl. **C.** Plots of the initial NADPH consumption rate (mM/minute) during the catalysis of GPx7 WT or GPx7 C86A as a function of PDI concentrations. Note that the rates were calculated by subtracting the rate of non-catalyzed reaction (No GPx7) from those of GPx7-catalyzed reactions (GPx7 WT or C86A).

### Preferential oxidation substrate of GPx7 among PDI family members

Previous studies by us and others suggest that PDI oxidases have high selectivity against the PDI-family members in the disulfide bond formation network (10,12,21,44,45). The present study indicated that whereas the PDI oxidation activity of GPx8 is minimal, GPx7 has significant PDI oxidation activity. To investigate GPx7 preference for the PDI-family members, five representative members of the PDI family (PDI, ERp46, P5, ERp57, ERp72), either independently or in a mixture of five, were subjected to oxidation by GPx7 in the presence of H_2_O_2_, and the time-dependent changes of their redox states were monitored by immunoblotting with an antibody against each PDI (Fig. 6A). The results of GPx7-catalyzed single-PDI oxidation demonstrated that the enzyme was a versatile oxidase to all PDI family members tested (Fig. 6A & B, left). By contrast, when GPx7 was reacted with the mixture of PDIs, ERp46 and ERp72 were oxidized in greatest amount compared to the other three PDIs (Fig. 6A & B, right), suggesting that GPx7 has significant preference for these two members. Interestingly, even though P5 seemed a good substrate of GPx7 in the single-PDI oxidation assay, P5 remained fully reduced in the mixture of PDIs, probably due to the competitive inhibition by the other PDI family members. Unlike P5, ERp46 and ERp72 were significantly oxidized by GPx7 even in presence of the other representative PDIs (Fig. 6A & B, right).

**FIGURE 6.**
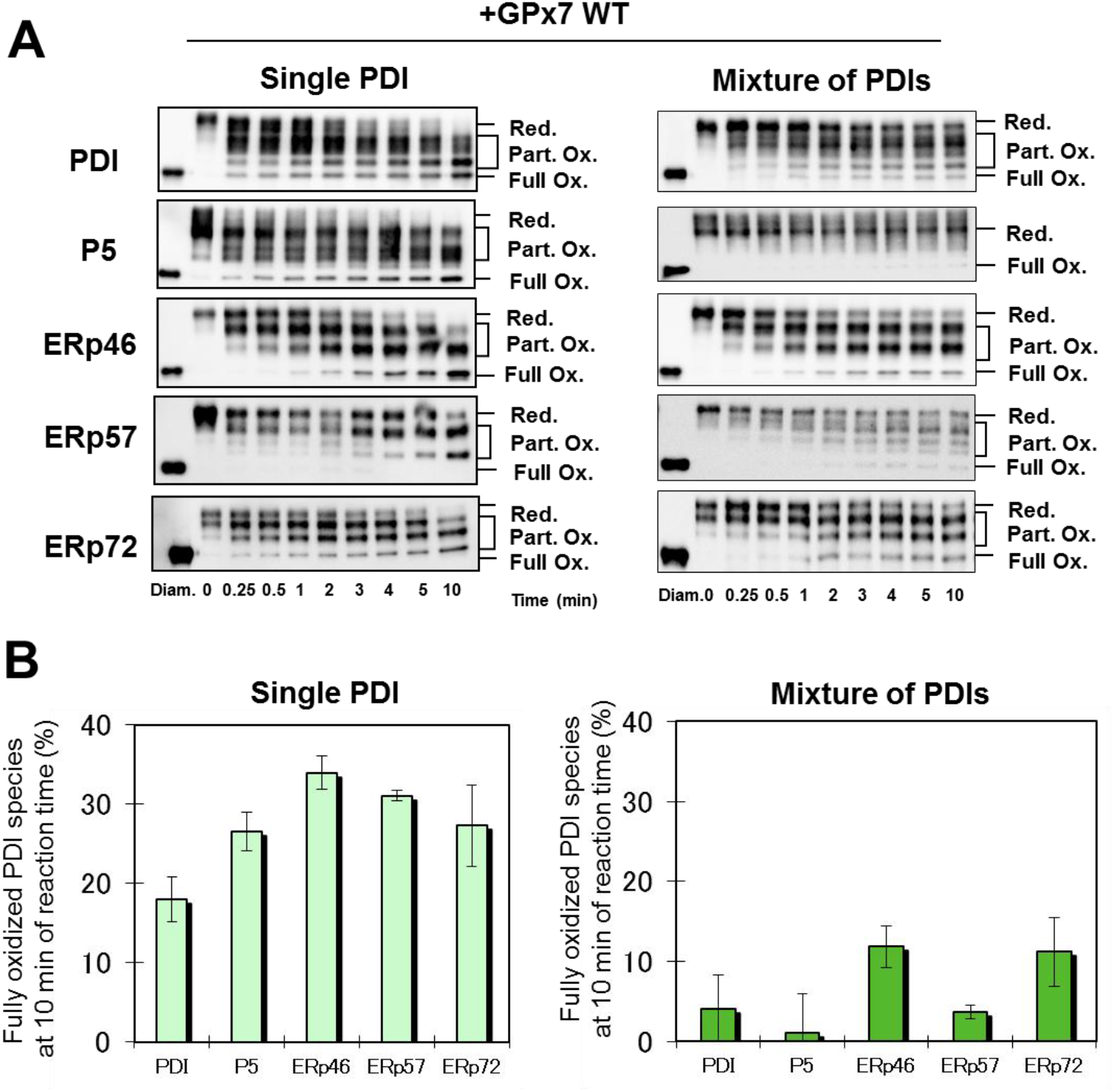
Preferential oxidation of PDI family members by GPx7. **A.** Time course of redox state changes of PDI-family members (PDI, P5, ERp46, ERp57 and ERp72) during the catalysis of PDI oxidation by GPx7 WT. Reactions were initiated by reacting 5 μM reduced PDIs (single or a subset of five) with 0.5 μM GPx7 WT in the presence of 50 μM H_2_O_2_. Reactions were quenched by cysteine alkylation with mal-PEG 2k at the indicated time points, followed by gel separation using non-reducing SDS-PAGE and monitored by immunoblotting with antibodies to each PDI family member. **B.** Quantification of the percentage of fully oxidized PDIs species generated after 10 minutes of reaction.

### H_2_O_2_-induced complex formation between PDIs and GPx7/8 in cells

To gain insights into physiological redox partners of GPx7 and GPx8, we next transfected HeLa cells with plasmids expressing FLAG-tagged Prx4, GPx7, or full-length GPx8, treated the cells with or without 5 mM H_2_O_2_ for 10 min, and alkylated the free cysteines with *N*-maleimide (NEM) as described in Experimental Procedures, followed by immunoprecipitation and immunoblotting with an antibody against each PDI family member (Fig. 7). We thus reproduced our previous observation that Prx4 most preferentially bound ERp46 and P5 in an H_2_O_2_-independent manner in cells (10). Notably, GPx7 bound PDI only slightly and did not bind the other PDI family members without H_2_O_2_ (Fig. 7). However, H_2_O_2_ treatment significantly increased amounts of PDI and P5 bound to GPx7. Similar observation was made with GPx8 despite even smaller amounts of PDI and P5 co-immunoprecipitated with GPx8 (Fig. 7). Thus, GPx7 displayed PDIs preference different from Prx4 although both the peroxidases bound their preferential PDIs more tightly in the presence of H_2_O_2_. Notably, the intracellular redox partners of GPx7 did not coincide with its preferential substrates identified by *in vitro* experiments (Fig. 6), suggesting the presence of mediator proteins that link GPx7 to its physiological redox partners (see also Discussion).

**FIGURE 7.**
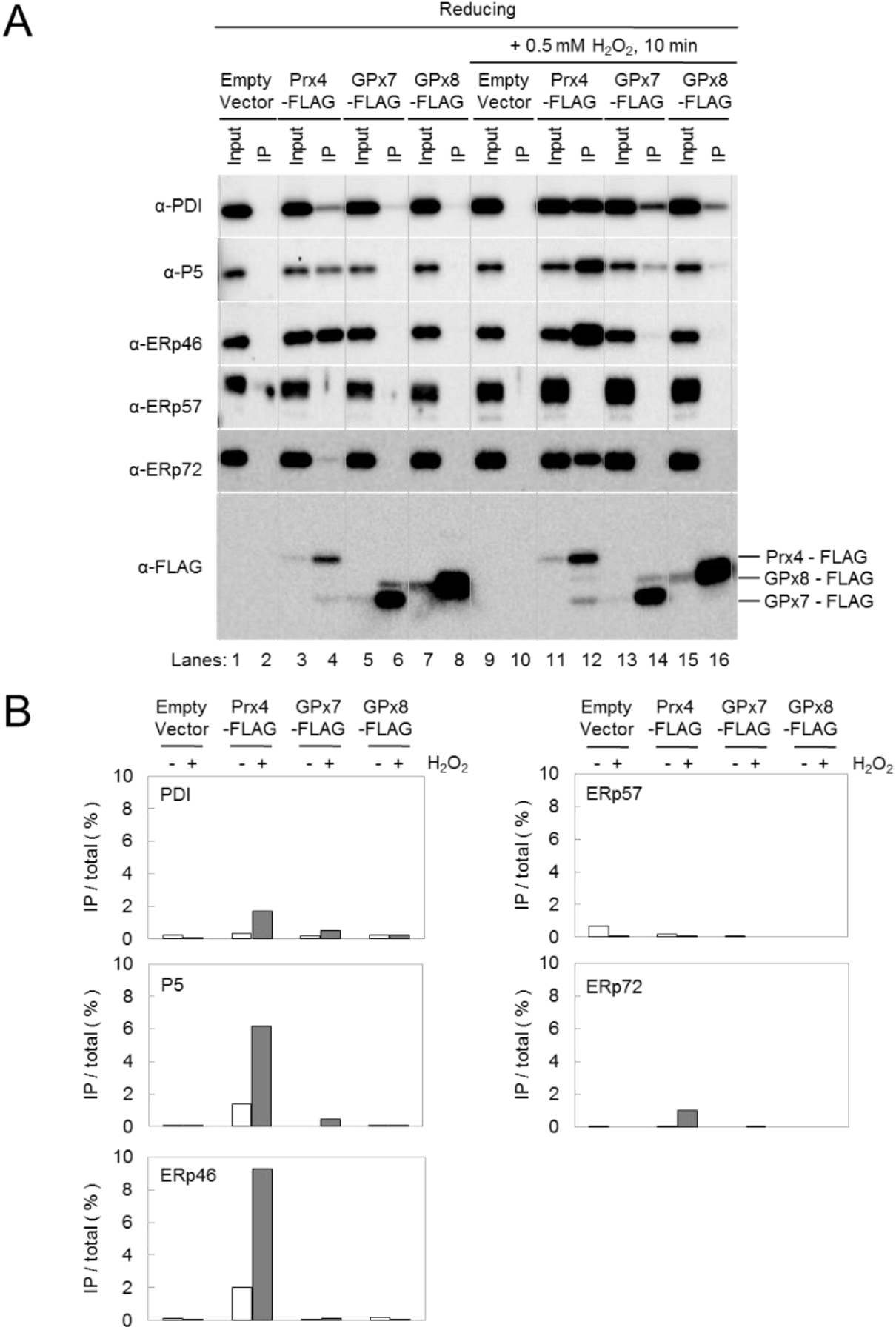
Complex formations of Prx4, GPx7 and GPx8 with PDI family members in cells. **A.** HeLa cells were transfected with pcDNA3.1^+^ (empty vector) (lanes 1, 2, 9, and 10), pCMV-Gg PRX4 (encoding PRX4-FLAG) (10) (lanes 3, 4, 11, and 12), pES104 (encoding GPx7-FLAG) (lanes 5, 6, 13, and 14), or pES105 (encoding full-length GPx8-FLAG) (lanes 7, 8, 15, and 16), grown in DMEM medium supplemented with 10% FBS for 24 h, treated with 0.5 mM H_2_O_2_ (lanes 9-16) for 10 min or not (lanes 1-8), washed twice with PBS, treated with 10% TCA (trichloroacetic acid), and subjected to alkylation with NEM (51). The NEM-treated cell lysates were subjected to immunoprecipitation with anti-FLAG M2 magnetic beads (Sigma) to purify complexes containing FLAG-tagged proteins. The immunoprecipitates were separated by reducing SDS-PAGE, and PDI family members and FLAG-tagged constructs contained in the immunoprecipitates were detected with antibodies to the indicated PDI family members or horse radish peroxidase-conjugated antibody to FLAG. Lanes 1, 3, 5, 7, 9, 11, 13, and 15 contained 1 μg of NEM-treated cell lysate (Input). Lanes 2, 4, 6, 8, 10, 12, 14, and 16 contained immunoprecipitates obtained from 50 μg of NEM-treated cell lysate. The positions of the FLAG-tagged constructs were indicated on the right in the bottom panel. **B.** The band intensities of PDI family members in the input and immunoprecipitate lanes on panel A were quantified using ImageJ 1.50i. PDI family members immunoprecipited with each FLAG-tagged peroxidase are shown as a percentage of total PDIs in the cell lysate.

## Discussion

Previously, GPx7 and GPx8 were demonstrated to be capable of utilizing Ero1α-derived H_2_O_2_ in cells (29). While GPx8 was reported to interact directly with Ero1α and scavenge the neighboring H_2_O_2_ molecule (39), the present work revealed that GPx7 had much higher reactivity with H_2_O_2_ than GPx8, leading to the more efficient PDI oxidation.

Reactivity of a given cysteine residue depends on the propensity of its thiolate form, which is quantitatively expressed as p*K*_a_ (46). However, p*K*_a_ is not the sole factor that determines the reactivity of a peroxidatically active cysteine with H_2_O_2_ (43,47). The active sites of GPx7 and GPx8 consist of a catalytic tetrad including Cys, Trp, Asn, and Gln/Ser residues. We here demonstrated that Gln92 residue in the catalytic tetrad of GPx7 is essential for the enhanced reactivity of GPx7 C_P_ with H_2_O_2_ probably because Gln92 serves to stabilize the sulfenylated C_P_ species via hydrogen bonding (Fig. 3 and Fig. 4B). Gln92 is highly conserved among all GPx family members but GPx8, where Gln is substituted by Ser114. In this regard, Ser114 can be interpreted to have an inhibitory role in the H_2_O_2_ scavenging activity of GPx8 (39), possibly providing GPx8 with other physiological functions in cells.

The present study also demonstrates that GPx7 WT preferentially employs a 2-Cys mechanism in oxidizing PDI while the GPx7 C86A mutant oxidizes PDI at a much higher rate via a 1-Cys mechanism. GPx7 C_P_ (Cys57) is oxidized by H_2_O_2_ to generate sulfenylated C_P_, which can in turn react with either reduced PDI or the C_R_ residue of GPx7 itself. The predominant usage of a 2-Cys mechanism by GPx7 suggests faster nucleophilic attack by the C_R_ residue of the same molecule than by the active-site cysteines of PDI (see Fig. 6A). Obviously, 2-Cys mechanism drives a much slower catalytic cycle than 1-Cys. This observation suggests that the intermolecular disulfide transfer from a Cys57-Cys86 pair of GPx7 to the PDI active sites is a rate-limiting step of the GPx7-mediated PDI oxidation cycle. In other words, the presence of a C_R_ residue serves to regulate the oxidative activity of GPx7. In this context, the selection of 1-Cys or 2-Cys mechanism by GPx7 could be an effective strategy to maintain the redox homeostasis in the ER, as is the similar case with Ero1α, which tightly modulates its oxidative activity through the disulfide bond rearrangement among the four regulatory cysteines in response to the redox environment in the ER (7–9,23). However, the higher concentrations of reduced PDI did not greatly enhance the oxidative activity of GPx7 WT (Fig. 5), making it unlikely that the switching from a 2-Cys to 1-Cys mechanism is induced in GPx7 when the ER environment becomes more reducing. It is to be further elucidated how GPx7 contributes in maintaining the ER redox homeostasis in concert with other redox partners.

In the previous study, Ero1α was shown to preferentially oxidize PDI, and to a lesser extent ERp46, whereas Prx4 preferential oxidation substrates were P5 and ERp46 (10,12,44). Here we demonstrated that whereas GPx7 preferentially oxidized ERp46 and ERp72 *in vitro,* the enzyme bound PDI and P5 in H_2_O_2_-treated cells. The complex formation between GPx7 and P5 in cells suggests that some third-party proteins may mediate the interaction between these two. In agreement with this notion, GPx7 was reported to form a disulfide-linked complex with BiP, an ER-resident chaperone under oxidative stress conditions (35), while P5 targets BiP client proteins (48). Moreover, the preceding work reported that large ER-localized multiprotein complexes including various molecular chaperones such as BiP, GRp94, PDI, ERdj3, Hsp40, cyclophilin B, ERp72, GRP170, UDP-glucosyltransferase, and SDF2-L1 are formed in the ER (49). GPx7 may be one of the yet identified components contained in this multiprotein complex although further extensive studies are required to clarify the diverse interaction network involving GPx7 in cells.

In conclusion, the present work demonstrates that whereas GPx7 is an efficient PDI peroxidase, its phylogenic cousin GPx8 does not possess an efficient catalytic activity. GPx7 and GPx8 are similar in overall structure, such that they could share the same H_2_O_2_-scavenging mechanism involving their catalytic cysteine pairs. However, the different local environments near the cysteines give rise to the strikingly different reactivity with H_2_O_2_ of these two peroxidases. Although these two enzymes appear to fulfill distinct physiological roles in cells, it will be interesting to see if GPx8 can function as a backup enzyme of GPx7 under some special situations such as a highly reducing redox environment or a highly H_2_O_2_ abundant condition in the ER.

## Experimental Procedures

### Plasmids

Plasmids for overexpression of GPx7 and the luminal domain of GPx8 in *Escherichia coli* (pKEHS780 and pVD54, respectively) (29) were kind gifts from Dr. Lloyd Ruddock (University of Oulu). GPx7 and GPx8 Cys-Ala mutants were constructed by using the QuikChange Site-Directed Mutagenesis Kit (Agilent Technologies).

Primers and templates used to construct following plasmids are listed in Table 1. Plasmid pES101 (encoding GPx7) was constructed by assembling three DNA fragments using the Gibson Assembly Kit (New England Biolabs). The DNA fragments used were a fragment amplified from pcDNA3.1^+^ (Invitrogen) with primers NeoRH1 and gpx7E1, a fragment amplified from pKEHS780 with primers gpx7E2 and gpx7E4, and a fragment amplified from pcDNA3.1^+^ with primers NeoRH2 and gpx7E3. Plasmid pES102 that expresses the luminal domain of GPx8 in the ER, and plasmid pES103 (encoding GPx8) were constructed by assembling fragments amplified using a template and a primer set indicated on Table 2. Plasmids pES104 (encoding GPx7-FLAG on pcDNA3.1^+^) and pES105 (encoding GPx8-FLAG on pcDNA3.1^+^) were constructed by inserting a triple FLAG sequence in front of the KDEL sequence of these proteins. They were constructed also by assembling fragments amplified using a template and a primer set indicated on Table 2.

**Table 1.**
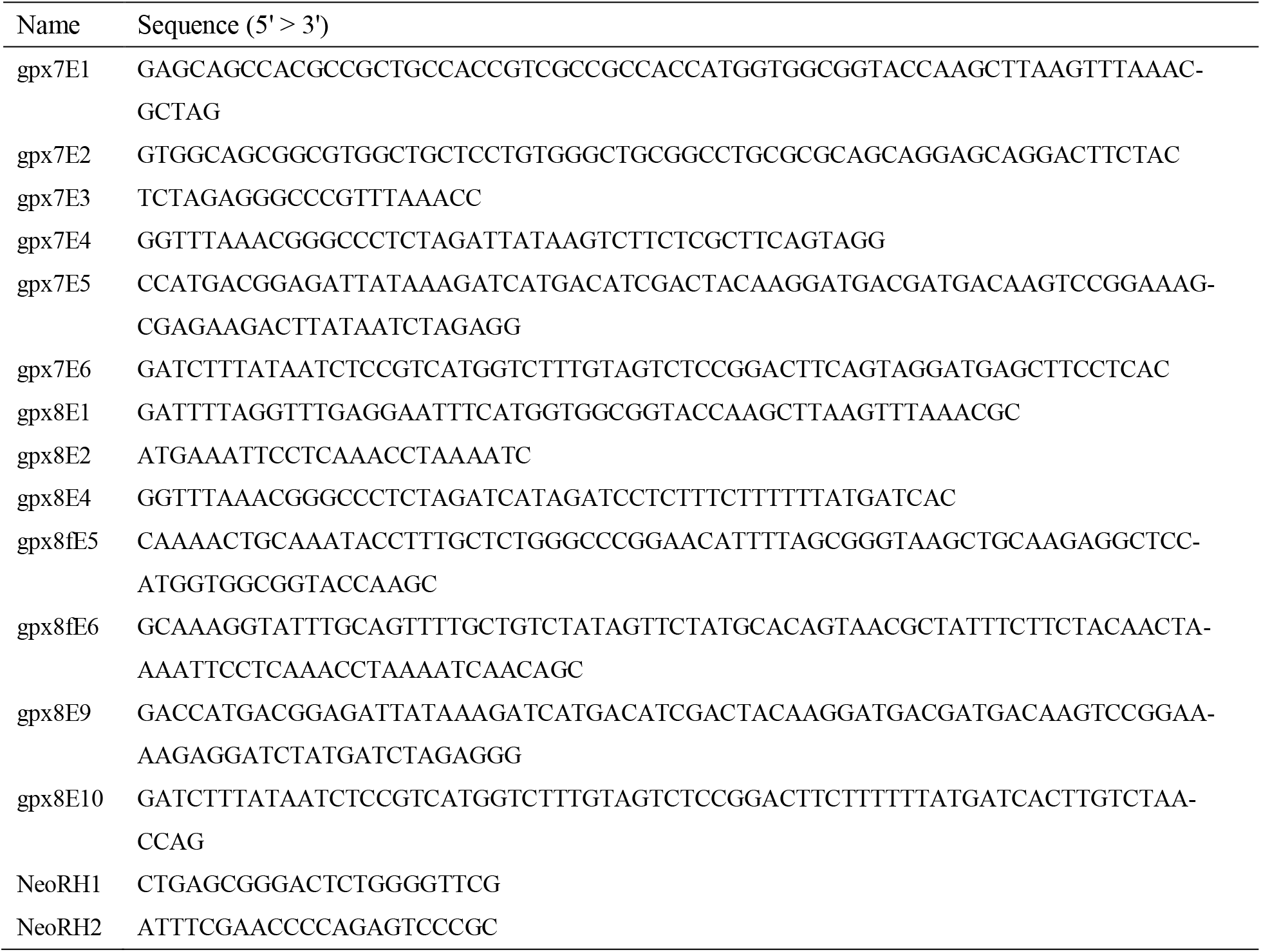
Primer sequences

**Table 2.**
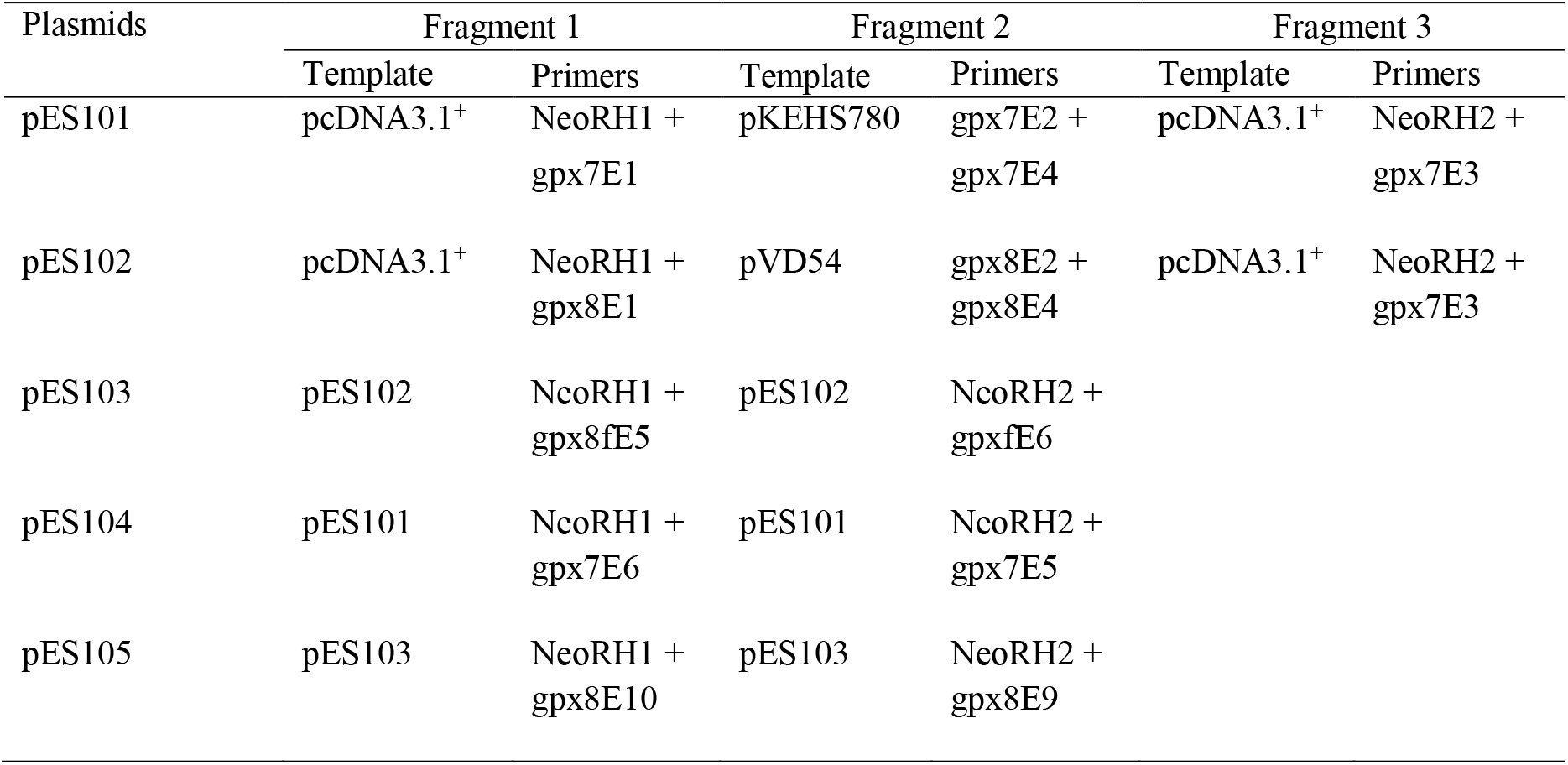
Template and primers used to construct pES plasmids

| Plasmids | Fragment 1 |  | Fragment 2 |  | Fragment 3 |  |
| --- | --- | --- | --- | --- | --- | --- |
|  | Template | Primers | Template | Primers | Template | Primers |
| pES101 | pcDNA3.1<sup>+</sup> | NeoRH1 +<br>gpx7E1 | pKEHS780 | gpx7E2 +<br>gpx7E4 | pcDNA3.1<sup>+</sup> | NeoRH2 +<br>gpx7E3 |
| pES102 | pcDNA3.1<sup>+</sup> | NeoRH1 +<br>gpx8E1 | pVD54 | gpx8E2 +<br>gpx8E4 | pcDNA3.1<sup>+</sup> | NeoRH2 +<br>gpx7E3 |
| pES103 | pES102 | NeoRH1 +<br>gpx8fE5 | pES102 | NeoRH2 +<br>gpxfE6 |  |  |
| pES104 | pES101 | NeoRH1 +<br>gpx7E6 | pES101 | NeoRH2 +<br>gpx7E5 |  |  |
| pES105 | pES103 | NeoRH1 +<br>gpx8E10 | pES103 | NeoRH2 +<br>gpx8E9 |  |  |

### Protein overexpression and purification

Plasmids for overexpression of GPx7, the luminal domain of GPx8, PDIs, and their mutants were transformed into *Escherichia coli* BL21(DE3). Cells harboring the plasmids were grown at 30°C and induced by the addition of 0.5 mM IPTG at an OD600 of 0.5. Cells were cultured for additional 4 h at 30°C and then harvested. To purify GPx7, GPx8, and their mutants, cells in buffer A (50 mM Tris/HCl, pH 8.1, 0.3 M NaCl, and 1 mM PMSF) were disrupted using a microfluidizer (Niro Soavi PA2K). After centrifugation of the cell lysate to remove cellular components at 12,000 rpm for 20 min, the supernatant was loaded onto an open Ni-NTA Sepharose column (QIAGEN). The column was washed with buffer A containing 20 mM imidazole, and proteins were eluted with buffer A containing 200 mM imidazole. The eluted sample was concentrated to 500 μl by filtration using Amicon Ultra filter units (MWCO, 10,000; Millipore), applied to a Superdex 75 size-exclusion column (GE Healthcare) pre-equilibrated with 50 mM Tris/HCl (pH 8.1), 0.3 M NaCl, and 1 mM EDTA, and finally eluted with the same buffer.

To purify PDIs, the supernatant of the cell lysate was applied to the open Ni-NTA Sepharose column. Fractions eluted with 200 mM imidazole were further purified by MonoQ anion exchange column (GE Healthcare) and Superdex 200 size-exclusion column (GE Healthcare).

### Antibodies, reagent, and immunological techniques

Rabbit antibodies to ERp57 (GTX100297), and to ERp72 (GTX115263) were purchased from GeneTex. For the detection of proteins fused with a triple FLAG tag, mouse anti-FLAG M2 antibody conjugated with horseradish peroxidase (Sigma A8592) was used. Rabbit antibody to PDI has been published (50). Antibodies to ERp46 and P5 were used as described previously (10). Immunoloblotting was performed using standard method. The protein bands were visualized using an appropriate secondary antibody, Clarity Western ECL Substrate (Bio-Rad) and ChemiDoc Touch Imaging System (Bio-Rad). The images were processed with Image Lab software (Bio-Rad).

### Analysis of the redox states of GPx7, GPx8, PDI and their mutants

Purified GPx7, the luminal domain of GPx8, PDIs and their mutants were separately reduced with 10 mM DTT for 10 min at 30°C followed by DTT removal through a PD-10 column (GE Healthcare) pre-equilibrated with 50 mM Tris/HCl (pH 7.4) buffer containing 300 mM NaCl. To analyze the redox status of GPx7, GPx8 and their mutants, different concentrations (10, 50 and 200 μM) of H_2_O_2_ was mixed with 1 μM reduced GPx7, GPx8 or their mutants in degassed buffer containing 50 mM Tris/HCl (pH 7.4) and 300 mM NaCl at 30°C. For PDI oxidation assays, 10 μM reduced PDI was incubated with 1 μM GPx7, GPx8 or their mutants, and reactions were initiated by adding 10, 50 or 200 μM of H_2_O_2_. At indicated time points, samples were quenched using 1 mM mal-PEG 2000 (NOF Corporation). The reaction mixture was boiled for 3 min after an addition of an equal volume of 2 × Laemmli buffer. All samples were run through non-reducing SDS-PAGE followed by staining with Coomassie Brilliant Blue (CBB) (Nacalai Tesque). The gel images were captured by using ChemiDoc Touch Imaging System (Bio-Rad).

### Oxidative folding assay of RNase A

RNase A from bovine pancreas (Sigma-Aldrich) was reduced and denatured with 100 mM DTT and 6 M guanidinium hydrochloride. The sample was passed through a PD-10 column (GE Healthcare) pre-equilibrated with 10 mM HCl aq. to remove the reducing and denaturing reagents. Reduced/denatured RNase A was diluted to 20 μM (~50-fold dilution) and incubated with 1 μM GPx7 or its mutant, and 10 μM PDI in a buffer (50 mM Tris/HCl, pH 7.4, and 300 mM NaCl) at 30°C, and reactions were initiated by adding 50 μM of H_2_O_2_. At indicated time points, samples were quenched using 1 mM 4-acetamide-4′-maleimidylstilbene-2,2′-disulfonic acid (AMS; Molecular. Probes, Inc.). All samples were separated by non-reducing SDS-PAGE followed by staining with CBB. The gel images were captured by ChemiDoc Touch Imaging System (Bio-Rad). The recovery of RNase A activity was measured by monitoring the linear increase in absorbance at 295 nm on a U-3900 spectrophotometer (HITACHI) at 30°C after addition of 1.25 mM cytidine 2′,3′-cyclic monophosphate monosodium salt (cCMP, Sigma-Aldrich) to reaction mixtures at indicated time points.

### NADPH consumption assay

NADPH consumption coupled with GPx7-catalyzed PDI oxidation was monitored by measuring absorbance change at 340 nm using a SH-9000 microplate reader (Corona Electric Co.). The solution was first prepared by mixing 1 μM GPx7 WT or GPx7 C86A, 5-50 μM reduced PDI, 1 mM GSH, 1 U glutathione reductase (GR) and 200 μM NADPH in a buffer (50 mM Tris/HCl, pH 7.4, and 300 mM NaCl), and reactions were initiated at 30°C by adding 50 μM of H_2_O_2_ to the solution. The initial NADPH consumption rates were calculated by considering the decrease of absorbance at 340 nm during the initial reaction time of 30 sec with a molar extinction coefficient value of 6200 M^−1^ cm^−1^ for NADPH.

### Preferential PDI oxidation by GPx7 *in vitro*

5 μM each of purified PDI family members (PDI, P5, ERp46, ERp57, ERp72) was mixed with 0.5 μM GPx7. Reactions were initiated by adding 50 μM of H_2_O_2_ to the solutions. At indicated time points, samples were subjected to 2 mM mal-PEG 2000 (NOF Corporation) followed by immunoblotting with the corresponding anti-PDI antibodies.

### Cell culture, transfection, and sample preparation

Cells were transfected with plasmids using Effectene (Qiagen). For the transfection, HeLa cells were plated in 10 cm dishes at 8×10^5^ cells/well, cultured for 24 h, and transfected with an appropriate plasmid, following manufacture’s instruction. The amount of plasmid DNA used for the transfection was 2 μg for pcDNA3.1^+^ and pCMV-Gg PRX4 (encoding PRX4-FLAG) (10), and 6 μg for pES104 (encoding GPx7-FLAG), and 6 μg for pES105 (encoding full-length GPx8-FLAG). The larger amount of plasmid DNA was used for transfection with the latter two plasmids because the levels of protein production from these plasmids were smaller than that from the former one when the same amount of plasmid DNA was used. At 24 h after the transfection, cells were treated with or without 0.5 mM H_2_O_2_ for 10 min, washed twice with PBS, treated with 10% TCA (trichloroacetic acid), and subjected to alkylation with NEM (*N*-ethylmaleimide) essentially as described (51).

### Immunoprecipitation

Five hundred μg NEM-treated cell lysate prepared as described above was diluted with ice-cold KI buffer [2% (w/v) Triton X-100, 50 mM Tris-HCl (pH 8.0), 150 mM NaCl, 1 mM EDTA] and centrifuged at 15,000 X g for 10 min at 4°C to obtain NEM-treated cleared cell lysate. To purify FLAG-tagged proteins (PRX4-FLAG, GPx7-FLAG, or GPx8-FLAG) from the cleared cell lysate, fifty μl of prewashed anti-FLAG M2 magnetic beads (Sigma) were incubated with the NEM-treated cleared cell lysate at 4°C for 3 h. The immune complexes were collected by magnetization and washed four times with 800 μl of ice-cold High Salt buffer [1% (w/v) Triton X-100, 50 mM Tris-HCl (pH 8.0), 1 M NaCl, 1 mM EDTA], and once with 800 μl of 10 mM Tris-HCl (pH 8.0). The immunoisolates were then released by incubating the sample at 37°C for 1 h with 65 μl of 2 X Laemmli sample buffer supplemented with 10 mM NEM, 2 μg/ml pepstatin A, 1 mM benzamidine, and 1 mM PMSF.

## Data availability

All data described are contained within the article.

## Acknowledgements

We are grateful to Dr. Lloyd Ruddock (University of Oulu) for providing plasmids for overexpression of GPx7 and GPx8 in *E.coli*.

## Abbreviations

ER: endoplasmic reticulum
PDI: protein disulfide isomerase
GPx7/8: glutathione peroxidase-7/8
Ero1α: ER oxidoreductin-1α
Prx4: peroxiredoxin-4
Trx: thioredoxin
GSH: reduced glutathione
Sec: selenocysteine
ROS: reactive oxygen species
C_P_: peroxidatic cysteine residue
C_R_: resolving cysteine residue
mal-PEG 2k: maleimidyl PEG-2000
NEM: *N*-maleimide
CBB: Coomassie Brilliant Blue
AMS: 4-acetamide-4 ′ -maleimidylstilbene-2,2 ′ -disulfonic acid
cCMP: cytidine 2 ′, 3 ′ -cyclic monophosphate monosodium salt
GR: glutathione reductase
TCA: trichloroacetic acid

